# MSMBuilder: Statistical Models for Biomolecular Dynamics

**DOI:** 10.1101/084020

**Authors:** Matthew P. Harrigan, Mohammad M. Sultan, Carlos X. Hernández, Brooke E. Husic, Peter Eastman, Christian R. Schwantes, Kyle A. Beauchamp, Robert T. McGibbon, Vijay S. Pande

## Abstract

MSMBuilder is a software package for building statistical models of high-dimensional time-series data. It is designed with a particular focus on the analysis of atomistic simulations of biomolecular dynamics such as protein folding and conformational change. MSMBuilder is named for its ability to construct Markov State Models (MSMs), a class of models that has gained favor among computational biophysicists. In addition to both well-established and newer MSM methods, the package includes complementary algorithms for understanding time-series data such as hidden Markov models (HMMs) and time-structure based independent component analysis (tICA). MSMBuilder boasts an easy to use command-line interface, as well as clear and consistent abstractions through its Python API (application programming interface). MSMBuilder is developed with careful consideration for compatibility with the broader machine-learning community by following the design of scikit-learn. The package is used primarily by practitioners of molecular dynamics but is just as applicable to other computational or experimental time-series measurements. http://msmbuilder.org

## I. INTRODUCTION

Molecular dynamics (MD) is a powerful probe into atomistic dynamics. Recent advances in technology (specialized hardware [1] or commodity GPUs [2]) and strategies (massively distributed architectures [3–5]) enable simulations to reach larger size and longer timescales. Increasing quantities of raw data require novel and sophisticated analysis techniques [6]. Markov state models (MSMs) have gained favor for drawing interpretable conclusions from time-series data [6–9]. Briefly, MSMs model dynamic systems using a set of discrete states and pairwise transition rates. From these models, the researcher can compute observables of interest and make predictions. These models are statistically rigorous and easy to interpret. Furthermore, MSMs are able to stitch together many independent simulation runs, allowing researchers to fully exploit distributed computing.

The idea of describing a system by its states and rates is natural for chemists and biologists, but the estimation of states and rates from finite data (perhaps molecular dynamics) is not obvious. From the introduction of MSMs to the biophysics community, algorithmic improvements for constructing MSMs and computing observables have been the focus of intense study. The practical implementation of these algorithms has spawned several historical packages for MSM construction [10–12]. Each of these packages was tied strongly to the best practices in MSM construction of the time. Due to the fast-moving research around MSMs, software re-writes were common [13, 14].

We introduce MSMBuilder 3, a community-driven, open source software package for constructing MSMs. MSMBuilder offers a curated selection of MSM construction algorithms based on modern advances in the field. MSMBuilder is implemented in the Python programming language with performance-critical components written in C. It exposes an extensible API modeled after that of scikit-learn. The modular design ensures MSMBuilder 3 is adaptable to future improvements in MSM construction. The package can be invoked directly from Python or via the command line.

Through two instructive examples, we showcase the capabilities of MSMBuilder. In the first, we use MSMBuilder to analyze a biological system of interest from a dataset composed of more than 20,000 trajectories. This example builds a single MSM using methods unavailable in previous tools. Due to rapid advances in MSM methods, a variety of modeling choices are now available to researchers. In the second example, we demonstrate how MSMBuilder’s implementation of scoring functionals can be used to choose among these methods.

## II. INSTRUCTIVE EXAMPLES

### A. Constructing an MSM

MSMBuilder allows rapid analysis of large molecular dynamics datasets. In this example, we construct an MSM of a kinase molecule. Kinases are critical enzymes that control cellular pathways. Malfunctions of kinases have been linked to many different cancers [15]. Here, we use MSMBuilder to study the c-Src kinase, a regulator of cellular growth [16], and demonstrate that the resulting MSM can capture activation dynamics. Understanding the activation process reveals atomistic, kinetic, and thermodynamic insights into the protein’s conformational heterogeneity, which can help design better therapeutics.

Broadly, the procedure for constructing an MSM is to define a set of states and then estimate transition rates among those states. Before beginning model construction, researchers must obtain time-series data they wish to model. Usually, this is the output of a molecular dynamics engine (MSMBuilder supports nearly every MD trajectory file format [17]), but it could also be experimental time-series measurements. For this example, we use a previously-generated MD dataset of the c-Src kinase, publicly available from the Stanford Digital Repository (SDR) ^1^.

The first step of model construction is to transform the raw Cartesian coordinates into vector features that are invariant to translation and rotation (fig. 1, step 1). Here, we project our trajectory frames onto the dihedral angles created by each set of four consecutive alpha carbons (α angles) [18]. This reduces the dimensionality of the data from 12,693 Cartesian coordinates to 518 features. The appropriate featurization depends on the particular system under study (see section II B). MSMBuilder offers a collection of featurization strategies with a unified interface. Popular features include backbone and side chain dihedrals (through the DihedralFeaturizer class), heavy atom or C_α_ contact distances (ContactFeaturizer), distance of reciprocal inter-atomic distances (DRIDFeaturizer) [19], and root mean squared deviation to a set of structures (RMSDFeaturizer). There are additional utilities for concatenation of multiple choices of features and feature scaling.

**FIG. 1.**
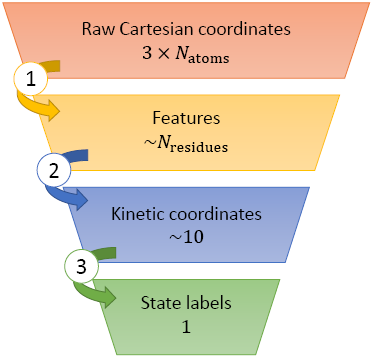
Data transformations and their dimensionality. Markov state models (MSMs) partition dynamical data into a set of states and estimate rates between them. A typical pipeline for state definition consists of a series of transformations (indexed by circled numbers) between representations of the data. Each step projects a higher dimensional representation onto a lower dimensional representation. The approximate dimension of each representation is reported below the representation name. Although not traditionally thought of as a dimensionality reduction, clustering (step 3) reduces each frame to a single integer cluster label.

The second step in MSM construction projects structural features onto a lower-dimensional subspace (fig. 1, step 2). This improves the statistical qualities of subsequent steps, but may discard important information if the projection is not carefully chosen. Time-structure based independent component analysis (tICA) finds a set of “slow” (high autocorrelation) coordinates. In practice, this dimensionality reduction has proven to be very useful for capturing slow, biophysical conformational change [20, 21]. In this example, we reduce the dimensionality of our kinase data from 518 dihedrals to 5 tICA coordinates. MSMBuilder includes support for similar algorithms (SparseTICA [22]) as well as general manifold learning algorithms like principal components analysis (PCA), SparsePCA, or MiniBatchSparsePCA. Prior to 2013, this step was not available for model construction. Accordingly, software available at the time could not easily be extended to accomodate tICA intermediate processing. The design of MSMBuilder 3 permits arbitrary addition, subtraction, and re-ordering of data transformation steps.

Next, we define the states of our MSM by grouping conformations which interconvert rapidly (fig. 1, step 3). For the c-Src kinase, we employ the MiniBatchKMeans [23] clustering algorithm to parition our data into 200 microstates. We note that our data has been reduced from 5 tICA coordinates to one integer cluster label per frame. The prior dimensionality reduction permits using off-the-shelf clustering algorithms. Accordingly, MSMBuilder supports K-Means like clustering algorithms (KCenters, KMedoids, and MiniBatchKMedoids), and hierarchical clustering.

With our states defined, we proceed to estimate the rates among them. As the final model construction step, we learn a continuous-time MSM [24] from our labeled trajectories. We have chosen to use a continuous-time MSM to directly estimate transition rates; we could have alternatively built a traditional MSM (to estimate transition probabilities) or a hidden Markov model (HMM). We direct interested readers to a more thorough application of HMM modelling to the c-Src dataset [25]. The relevant Python code for constructing this MSM is shown in fig. 2. Complete, executable code is available in the SI as an IPython [26] notebook.

**FIG. 2.**
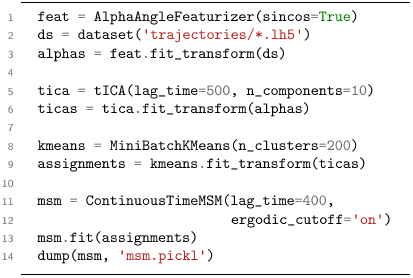
Sample MSM code. MSMBuilder balances a powerful API (application programming interface) with ease of use. A sample workflow is shown here using the Python API. Following the successful model of the broadly-applicable `scikit-learn` package, each modelling step is represented by an estimator object which operates on the data. Here, the `AlphaAngleFeaturizer` transforms raw coordinates into *α* angles. The output of this transformation is fed into the `tICA` dimensionality reduction, `MiniBatchKMeans` clustering algorithm, and finally into the `ContinuousTimeMSM` model. MSMBuilder provides a litany of utility functions for dealing with large molecular dynamics datasets for I/O. While this example shows the Python API, MSMBuilder is fully functional from the command line with an intuitive 1-to-1 correspondence between Python estimator objects and command-line commands.

To draw interpretable conclusions from our data via Markov modelling, we query the model. For c-Src, we use MSMBuilder to relate model behavior to biological function. We present a log-scaled 2D histogram (fig. 3a) of the trajectories projected onto the two dominant slow processes, or “tICs”, from our tICA model. We then sample the centroids of states (shown as pink and black stars) in low free energy regions to visualize representative configurations in three dimensions [27] (fig. 3c and d). The dominant tIC (x-axis) highly correlates with the activation of the kinase. Kinase activation requires the unfolding of the activation loop (red) and an inward swing of the catalytic helix (C-helix). The inward rotation of the helix coincides with the switching of hydrogen bonding pair from Glu-Arg to Glu-Lys (licorice). We investigate the dynamics between the active, inactive, and intermediate macrostates by applying Robust Perron Clustering Analysis (PCCA+) to our MSM. PCCA+ is a spectral clustering method, which lumps MSM states into an arbitrary number of metastable macrostates, facilitating qualitative analysis of rates and populations among biologically-relevant macrostates [28]. The rates among three macrostates are shown by the thickness of arrows in fig. 3b. Further options of querying the model (not shown here but available in MSMBuilder) include computation of relaxation timescales, transition path theory analysis [29–31], and generation of synthetic trajectories for visual inspection.

**FIG. 3.**
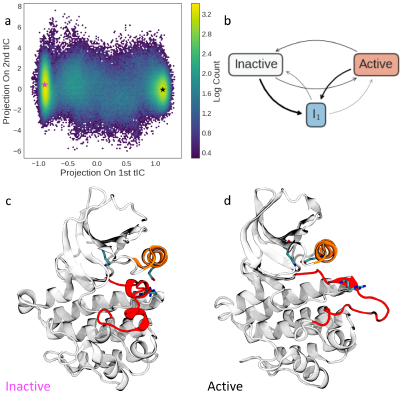
c-Src kinase MSM. MSMBuilder constructs interpretable models from large datasets. This figure shows a 2D-histogram for the Src kinase from tICA-MSM analysis projected onto the dominant modes of a tICA model (a). A simple macrostate model of the dynamics shows the presence of an intermediate state I_1_ connecting the inactive and active states (b). The arrow thickness corresponds to the rate of transitions. The model indicates that the active state (red) is the most stable state followed by the inactive and intermediate states (gray and blue, resp.). The analysis discovers a coordinate (the first tIC) between the known active and inactive conformations. Representative structures are selected from MSM states and show the conformational differences between the two basins. The unfolding of the activation loop (red helix) forms a catalytically active Src capable of initiating and regulating downstream signaling pathways (c and d).

The assortment of modeling options such as the choice of featurizer, the use of dimensionality reduction, and the selection of the clustering algorithm, along with any associated internal parameter choices, presents the modeler with a motley of modeling decisions and tunable parameters. In the next section, we show how a scoring metric for MSMs can provide the modeler with a unbiased protocol for determining which parameters are suitable given a set of MD trajectories.

Historically, the heuristic choice of hyperparameters—choices of protocol—rendered MSM construction as much of an art as a science. It is clear from section IIA that there is an abundance of algorithms available in MSMBuilder. In this instructive example, we use a scoring functional to select the best models.

Nüske *et al.* [32] introduced a variational principle that formalized the definition of a “good” MSM. In keeping with inspiration from the broader machine learning community, MSMBuilder extends this formalism in the context of cross-validation through the work of McGibbon and Pande [33]. The resulting generalized matrix Rayleigh quotient (GMRQ) score offers an objective way to pick the best model (i.e. the appropriate modeling choices) from the given data. Briefly, the GMRQ measures the ability of a model to capture the slowest dynamics of a system. The variational principle states that approximating the full phase space by discrete states will always yield dynamics that are too fast. The GMRQ score is a summation of the leading eigenvalues of the model and therefore provides a measure of “slow-ness”. A higher score means the model is closer to the variational bound, and therefore should be prefered over lower scoring models.

In this example, we use the GMRQ score under cross-validation to evaluate the relative merit of enumerated hyperparameter values when constructing a model for the Fs peptide [34]. The relevant code in fig. 4 sets up a choice between two structural features (dihedral angles or contact distances) and a choice among tICA lag times. We perform shuffle-split cross-validation by randomly assigning the 28 trajectories to either the training set or test set. The MSM is learned on the training set and scored on the test set. By concealing the training data during scoring, cross-validation guards against overfitting (overconfidence in excessively complex models). The trajectories are re-shuffled and this process is repeated to compute an average score for a given set of hyperparameters. The scores for each of the 50 cross-validation splits are plotted in a box plot in fig. 5. The dihedral angle featurization with a lag-time of 4 steps gives the best model in this search space. A simple grid search as performed in this example can become intractible as the number of hyperparameters (i.e. the dimension of the search space) increases. We direct interested users to Osprey [35], a tool for hyperparameter optimization with a variety of search strategies and support for parallel computation. Osprey interoperates with any scikit-learn estimator including those in MSMBuilder.

**FIG. 4.**
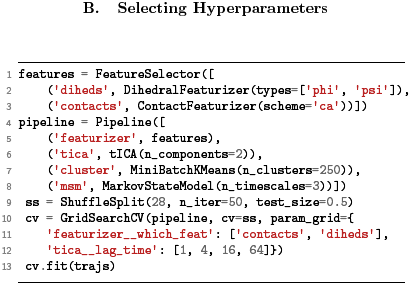
Sample GMRQ code. MSMBuilder seamlessly interoperates with the broader Python ecosystem. In this code sample, we use scikit-learn for algorithm-agnostic data processing and MSMBuilder for biophyics-oriented time-series algorithms with the goal of selecting model hyperparameters. Our analysis pipeline is similar to that of section IIA but with a choice of features (between dihedrals and contact distances) and tica lag times (among 1, 4, 16, and 64 steps). The ShuffleSplit cross-validation scheme runs 50 iterations of equal partitioning of the 28 trajectories between train and test sets, and we perform a full grid-search over parameter choices. We can plot the distribution of scores vs. parameters as in fig. 5.

**FIG. 5.**
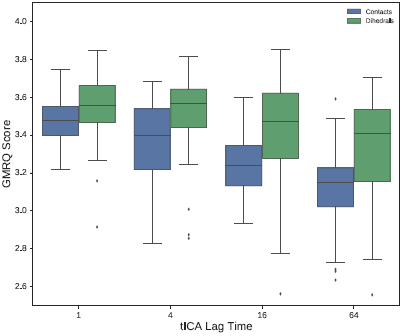
GMRQ parameter selection. MSMBuilder offers robust machinery for selecting hyperparameters that cannot *a priori* be learned from the data. Here, we perform shuffle-split cross validation over choices of featurization and tICA lag time. Historically, these parameters were chosen heuristic ally. With the advent of the GMRQ score and its implementation in MSMBuidler, we can choose these parameters in a statistically rigorous way. Here, we plot the distribution of scores for each set of of model parameters. Note that a higher score is generally an indication of a more predictive model. In this example, we find that featurization with dihedral angles at a lag time of 4 steps has highest median score and recommend this hyperparameter set to be chosen for the final model.

This example leverages the Pipeline, ShuffleSplit, and GridSearchCV machinery from scikit-learn. Additionally, MSMBuilder uses this library internally for generic machine learning algorithms such as clustering or PCA. We note that such general algorithms do not need to be reinvented and re-programmed by the biophysics community. By delegating some development effort to this widely-used machine learning library, we ensure that the development of MSMBuilder is focused on biophysical algorithms and considerations. This advantage offers rapid adoption of the latest algorithms which have demonstrated improved ability to build MSMs (e.g. [33]) and a larger community for code maintenance and longevity.

## III. CONCLUSIONS

MSMBuilder 3 is a powerful and accessible software package for drawing interpretable conclusions from time-series data. We used two examples to demonstrate how MSMBuilder can make sense of a molecular dynamics dataset consisting of thousands of trajectories in a highly automated and statistically robust way. In the first example, we construct a “vanilla” MSM and show how MSMBuilder enables the construction of interpretable models that expose the connection between biological function and structural dynamics. We highlight the breadth of relevant algorithmic choices for featurization, normalization, dimensionality reduction, clustering, and MSM modelling. In the second instructive example, we acknowledge that the explosion of choices in parameters and protocol can be overwhelming. We use the GMRQ score and off-the-shelf cross-validation machinery to do a simple grid search over tunable parameters to evaluate the relative merit of many MSMs built on the same MD dataset of a small protein. Since cross-validation is not a technique unique to biophysics, we leverage the greater Python machine learning ecosystem for this example.

More broadly, MSMBuilder’s power and clarity is derived from its integration with the machine learning community at large. Our power to focus on developing methods bespoke to biophysics and time-series analysis comes from exploiting general-purpose algorithms implemented by respective experts. The clarity of MSMBuilder’s API is due in large part to the massive amount of effort and skill put into the design of scikit-learn’s API. As distributed computing and Markov modelling continue to become more prominent, MSMBuilder offers a sustainable, extensible, powerful, and easy-to-use set of Python and command-line tools to help researchers draw meaningful conclusions from their data.

## IV. AVAILIBILITY

MSMBuilder documentation and installation is available at http://msmbuilder.org. The source code is available under the open-source LGPL2.1 license and is accessible at http://github.com/msmbuilder/msmbuilder. The current release at time of writing is version 3.5 [36]. Complete examples can be found as IPython notebooks in the supporting information and at http://github.com/msmbuilder/paper.

## V. AUTHOR CONTRIBUTIONS

MPH, MMS, CXH, and BEH wrote the paper. MPH, MMS, CXH, BEH, PE, CRS, KAB, RTM, and VSP edited the paper. MPH, MMS, CXH, BEH, PE, CRS, KAB, and RTM wrote the software. VSP supervised the project.

## VI. ACKNOWLEDGMENTS

We extend thanks to all our contributors including Stephen Liu, Patrick Riley, Steven Kearnes, Joshua Adelman, and Gert Kiss. We acknowledge funding from NIH grants U19 AI109662 and 2R01GM062868. MMS acknowledges support from NSF-MCB-0954714. CXH acknowledges support from NSF GRFP (DGE-114747). KAB acknowledges support from NIH grant P30CA008747, the Sloan Kettering Institute, and Starr Foundation grant I8-A8-058. We acknowledge members of the Chodera, Pande, and Noë labs for helpful discussions. We thank Ariana Peck for invaluable feedback on this manuscript.

## Footnotes

1 Available here: https://goo.gl/LLchMT. For simulation details, see Shukla *et al.* [16].

